# Visual Attention doesn’t Travel, it Teleports

**DOI:** 10.64898/2026.08.07.743495

**Authors:** Kabir Arora, Surya Gayet, J. Leon Kenemans, Marnix Naber, Samson Chota, Stefan Van der Stigchel

## Abstract

Attention forms a key, yet elusive component of visual processing. We shift attention constantly across the visual field to enhance processing of relevant locations or stimuli in service of goal-directed behavior. Here, we investigate a fundamental property of attentional shifts: when covertly shifting from one location to another, does visual attention “travel” (enhance processing at intermediate locations) or “teleport” (not interact with intermediate locations)? While most shifts of information or movement “travel” (e.g., eye and body movements, neuronal signalling), for attentional shifts such intermediate processing enhancement might not be necessary, nor functional. To answer this question while tackling the difficulty of tracking covert attentional dynamics, we conducted an EEG-eyetracking experiment (n=24) paired with Rapid Invisible Frequency Tagging (RIFT). This let us track attentional enhancement during covert attentional shifts with high temporal and spatial precision. We successfully registered covert shifts, but found no evidence for any attentional modulation in-between the start and end-point of an attentional shift. Additional insilico modelling of attentional shifts confirmed that our method was sensitive enough to pick up on these modulations had they been present. Our results support a “teleporting” model of attention, suggesting that attention is implemented in a fundamentally different manner compared to overt visual behaviours such as eye movements.

## Introduction

Attention forms a fundamental component of visual processing. Within the complex visual world, not every piece of information is equally valuable at a given point of time. Only specific locations tend to be most relevant, and attention allows us to prioritize these by enhancing visual processing there. The location that has the most relevant information changes over time, and the ability of attention to then flexibly shift its focus towards other locations helps us to make sense of what we see. The neural and behavioral signatures of attentional shifts are well-documented (Ben Hamed, 2025; Carrasco, 2011; Posner, 1980). However, we still know relatively little about how this shifting is actually carried out: when attention shifts from one location to the other, does it “travel” or “teleport”? That is, does it enhance visual processing at intermediate locations between its start and end point (“travel”), or, does it only enhance processing at the desired endpoint without interacting with intermediate locations (“teleport”)?

An initial, intuitive answer to this question, based upon familiar bodily movements, might be that attention travels through space when shifting from one point to another. This is because essentially any function that the body performs involves travelling from one point to the other, whether it is motor output (body parts must travel through 3D space to reach a particular endpoint), or neuronal signaling (information is passed through the nervous system continuously as travelling changes in potential). Even within the field of vision, the dominant phenomena of eye movements works in this manner: gaze position simply must travel through all intermediate points in visual space when making eye movements between two locations. On the other hand, attention is not bound by the same physical rules as other bodily movements; i.e., it does not need to travel through all intermediate locations in the visual field when shifting position. Attentional facilitation implemented through, for example, an enhancement in visual processing, can simply end at one location, and begin at another. It often serves no functional purpose to contiguously attend all locations between a desired start and endpoint, especially if actively attending requires effort. This might simply be too costly a process to be worth implementing. It is thus also fair to expect that during an attentional shift, attention might simply “teleport”. That is, shifting attention might only involve changes in enhancement at the attentional point of origin and arrival, not at intermediate points.

Previous lines of research have asked similar questions about the manner in which attention traverses visual space. Attention has been observed to travel across the visual field, but this has mainly been in tasks where the intermediate space between start and endpoint was still in some way task relevant, such as stimulus-tracking (Shih & Sperling, 2002; Shioiri et al., 2002), curve-tracing (Houtkamp et al., 2003; Scholte et al., 2001), or object-based attention (Jeurissen et al., 2016; Pooresmaeili & Roelfsema, 2014). Investigating whether the reallocation of spatial attention fundamentally operates by traveling or teleporting thus necessitates a situation where the intermediate space is not functionally relevant (as is arguably the case during most shifts of attention that we perform throughout the day).

This poses a methodological challenge: dynamically tracking attention across ‘empty’ space in real time is notoriously difficult; it is, after all, “covert”. To the best of our knowledge, very little work has been able to record attentional shifts to directly address what occurs along its trajectory. A recent study used pupil-size modulations evoked by task-irrelevant probes between attended locations, and suggested that attention indeed travels throughout the spatial interval (Naber et al., 2026). Behavioural responses to similar task-irrelevant probes between attended locations have also suggested a travelling model of attention (Shulman et al., 1979). However, even such targeted work contains an inherent limitation: even though the probes presented here in the intermediate locations were task-irrelevant, they were still visible. The limitation here is that the observed attentional effects may have been caused by introducing such visible probes that would not have been present during ‘natural’ shifts of attention.

A recent technique offers a promising solution for the complex problem of concurrently tracking attentional allocation at several locations with high temporal resolution. Rapid Invisible Frequency Tagging (RIFT) (Arora, Hustá, et al., 2026; Zhigalov et al., 2019) uses imperceptibly flickering stimuli to uniquely track visual processing (and attentional modulations to this processing) at various locations across the screen. RIFT stimuli are invisible. This means that RIFT goes beyond the traditional SSVEP approaches of studying attention (Müller et al., 1998; Norcia et al., 2015), since it does not display any visible probes that might interfere with attentional allocation. Here, we use six simultaneous RIFT tags to track the spatio-temporal dynamics of an attentional shift. We find no enhancement of visual processing during an attentional shift in between its start and end locations, suggesting that visual attention “teleports”, that is, does not facilitate the processing of intermediate locations. We also conducted additional in-silico modelling of attentional shifts to confirm that our method was sensitive enough to pick up on these modulations if they had been present. More broadly, this study forms a first step towards non-invasively tracking attention over the whole visual field to reveal insights in the various domains of visual cognition where attention plays a role.

## Results

To investigate how covert attention moves across the visual field, we ran an EEG-Eyetracking experiment paired with Rapid Invisible Frequency Tagging (RIFT) in 24 healthy human participants. Participants were shown a propeller-shaped stimulus with four segmented wedges (Figure 1A). A spatial, arrow-shaped cue appeared at fixation, and cued one of its 4 outer segments (left/right/up/down) with 100% validity. At the corresponding outer segment, participants were presented with a Landolt-C target and performed a simple orientation discrimination task. All participants performed the task with an accuracy (mean 72.5%, 95% bootstrapped CI: [72.0%, 73.1%]) that was close to the staircased level (70%), and significantly above chance (25%).

**Figure 1:**
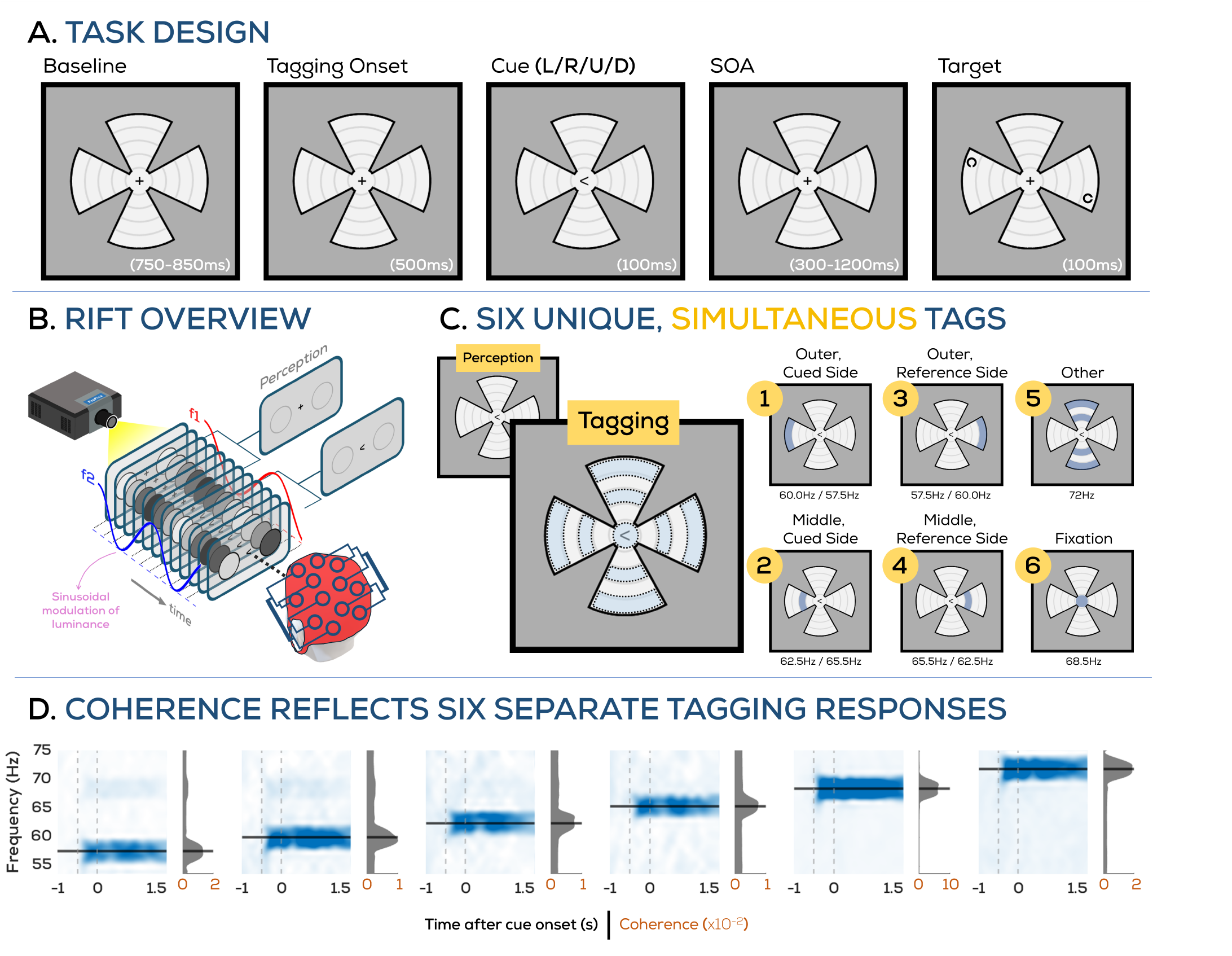
Task design and RIFT implementation. **A. Task:** Participants were presented with a propeller-shaped display, and reported the orientation of a landolt-C stimulus, which was preceded by a spatial cue indicating its upcoming location (left/right/up/down). Our task aimed to evoke covert shifts of attention from fixation to the outermost section of the propeller-shaped stimulus. **B. RIFT:** We implemented Rapid Invisible Frequency Tagging, a technique where parts of the display are flickered beyond the threshold of perception. The neural responses to these flickers are later recovered from the EEG to reveal attentional allocation across various tagged locations or stimuli. **C. Tagging during the task:** Here, we used 6 unique tagging frequencies to tag nine patches within the display; specifically, an outer (1) and middle (2) tag on the cued side, an outer (3) and middle (4) tag on the reference (opposite) side, the other middle and outer patches (5), and a circle around fixation (6). **D. All tagging responses were successfully recovered from the EEG**. Spectrograms show mean coherence across participants and top selected channels (see *“RIFT Response: Coherence”* in Methods) over the duration of the trial. Each of the six spectrograms shows the coherence response when phase-locking to the corresponding frequency tag (see *“Phase-skewing approach”* in Methods). Dotted vertical lines reflect tagging onset and cue onset.

Attentional allocation was measured at several different locations using imperceptible RIFT tags (Figure 1B-C). First, we verified that we could successfully recover all of these tagging signatures from the EEG data (Figure 1D). Coherence, which is a measure of the strength and phase-locking of the RIFT response across trials (see *“RIFT Response: Coherence”* in Methods), showed clear spectral peaks at each of the six frequencies (57.6Hz, 60.0Hz, 62.5Hz, 65.5Hz, 68.6Hz, 72.0Hz) that were used in the task.

On each trial, only one of the four wedges of the propeller-shaped stimulus (and specifically, only the outermost segment of this wedge) was cued. By tagging fixation, the most peripheral section (the “outer” tag), and the location between fixation and this outer tag (the “middle” tag), we aimed to measure whether visual processing, as measured by RIFT coherence, was enhanced at intermediate locations during an attentional shift. For both the outer tag and the middle tag, we compared the strength of the RIFT response elicited from the tags along the direction of the cued location to those in the opposite direction (the “reference” tags; these were automatically balanced in terms of eccentricity and tagged area, but not cued). First, we looked at the cued location, i.e., the “outer” tags (Figure 2A). As expected, RIFT responses were stronger at the cued location as compared to the outer tag on the reference side (significant cluster from 0.39s-1.18s after cue onset, p < 0.0005). This confirms that participants indeed covertly shifted their attention to the cued location. Next we assessed our central question of whether attention facilitated the processing of intermediate locations while shifting. We assessed this by comparing coherence at the middle tag in the cued direction to the middle tag on the reference side (Figure 2B). This comparison showed no attentional facilitation at the middle location prior to that already observed at the outer location. There was, however, attentional facilitation at this middle location later in the trial (two significant clusters; 0.58s-0.92s and 0.97-1.17s after cue onset, p = 0.0005 and p = 0.046 respectively).

**Figure 2:**
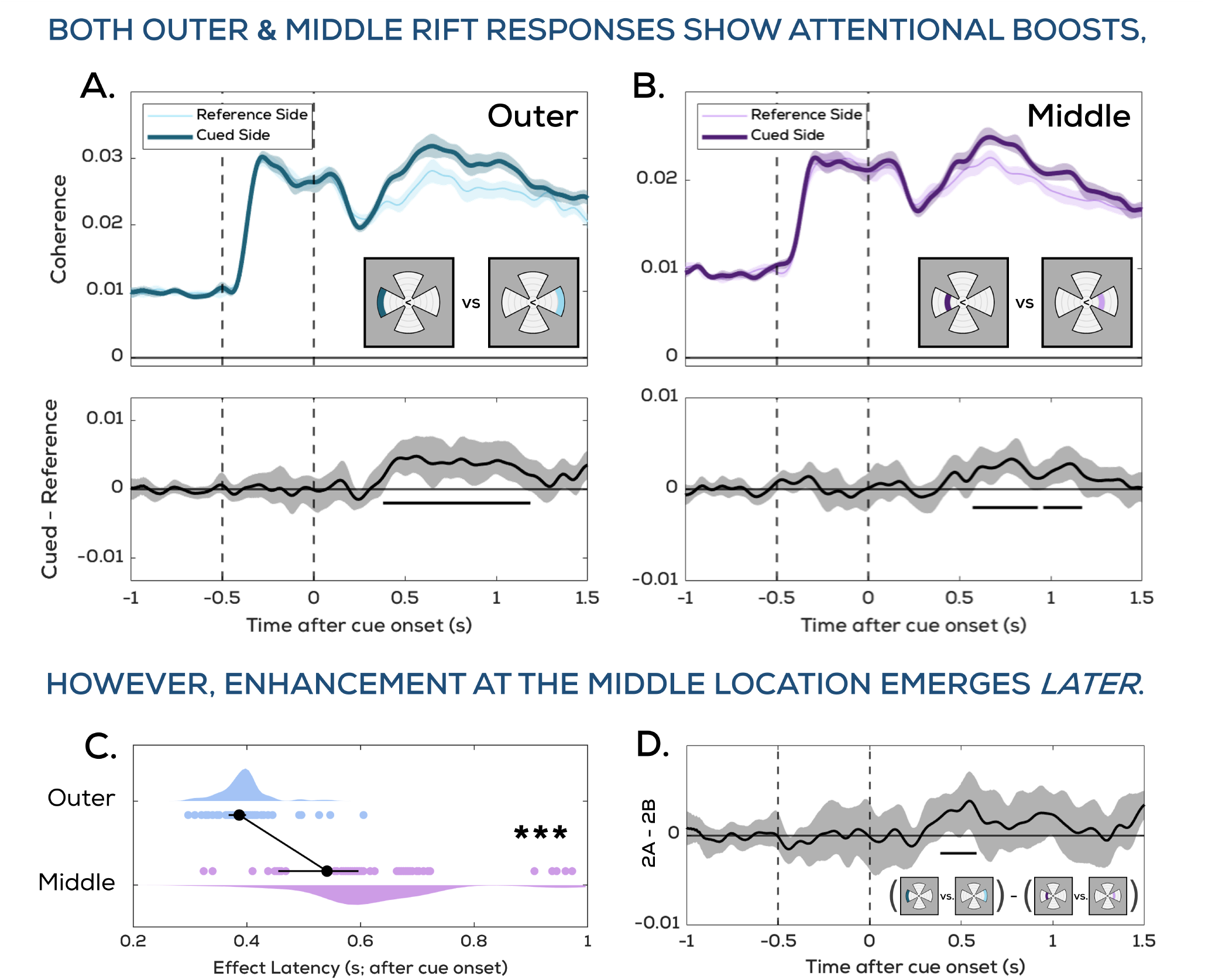
Attentional modulation of RIFT coherence of tags in the cued direction. **A**. RIFT responses are significantly stronger from the cued location (outer tag on the cued side) as compared to the outer tag on the reference side. **B**. RIFT responses are also significantly stronger from the middle location on the cued than the reference side, however, this difference does not emerge until later in the trial after the cued location is already boosted. This is confirmed by **C**. testing onset latency differences (see *“Latency comparisons”* in Methods for details), which show a significantly earlier effect onset for the outer than the middle location, and by **D**. testing the difference of differences traces from A and B, which shows a positive deflection, indicating that the outer location effect was stronger before the middle location effect. Traces in **A-B** reflect time-varying coherence values, black traces in **A-B** and **D** reflect difference traces, and task panels show which regions are being compared. All shaded regions around traces reflect 95% boostrapped CIs (cousineau-corrected), and horizontal bars indicate significance as per cluster test. Shaded regions in C reflect kernel density estimates of the corresponding distributions.

While we primarily asked here how shifts of attention impact intermediate locations between the shifts, our tagging display could also be used to assess how attention is divided over the whole stimulus display. This may reveal, for example, whether attentional enhancements at specific locations require suppression of visual processing at other, irrelevant locations. Or, it may reveal the speed of the attentional shift. We do so here not with the goal of producing a global estimate for how long it generally takes the visual system to shift attention, but instead, to get an estimate of shift duration from our data as a starting point for a later simulation analysis that determined whether our RIFT measure was sensitive enough to show an attentional modulation if attention indeed had “travelled” for the predicted duration. To assess how attentional allocation over the whole stimulus display evolved over time (as compared to the reference side), and attempt to pick up on attention “leaving” fixation, we looked at differences in coherence amplitudes relative to the tagging baseline periods for each tag (Figure 3; Supplementary Video 1).

**Figure 3:**
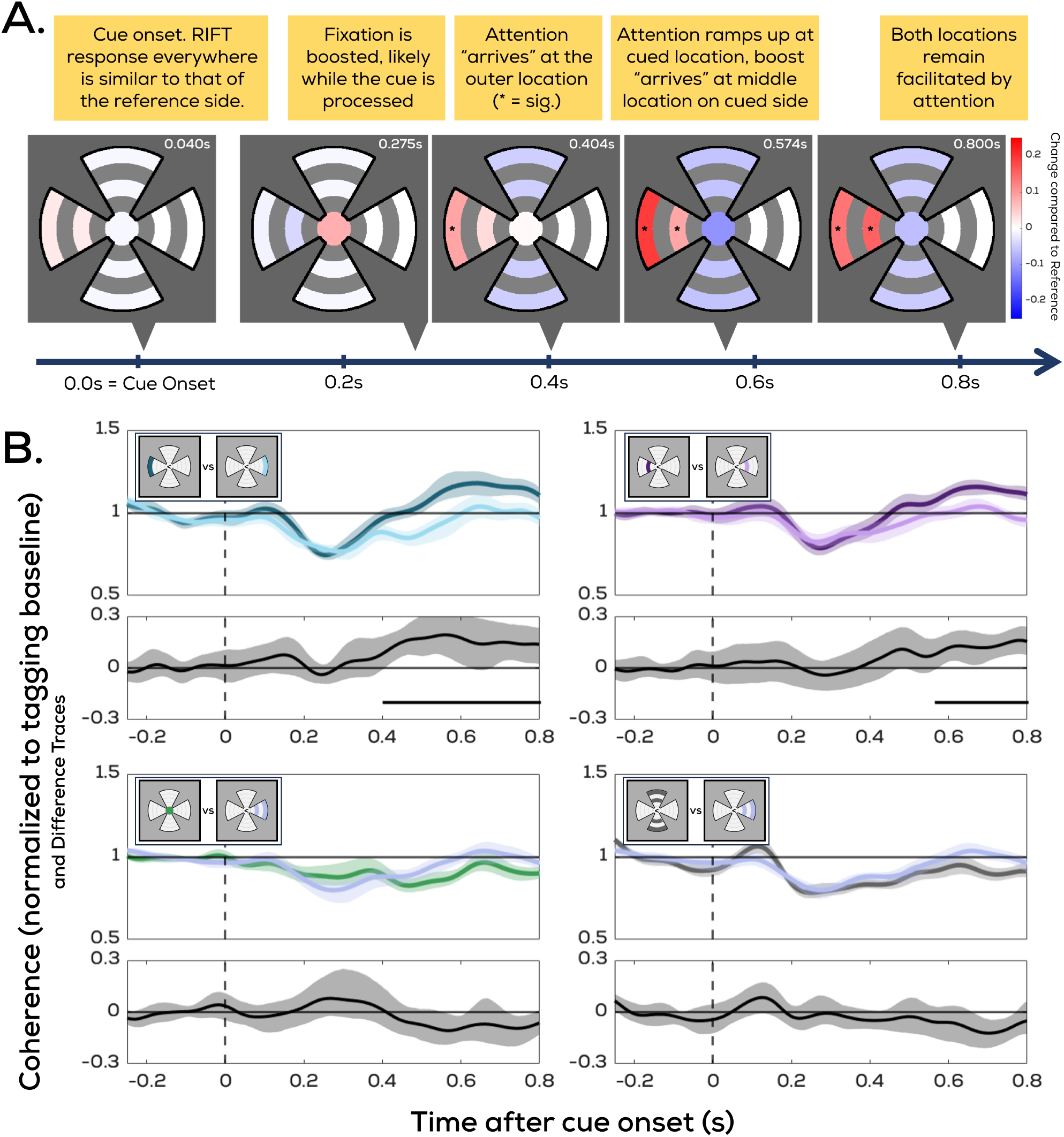
Attentional allocation over time across the visual field during a spatial shift of visual attention. **A**. Spatial visualization of noteworthy moments from traces in **B**. Pairwise comparisons of specific tags (cued location, cued side middle, fixation, and other) with corresponding patches on the reference side. Fixation and other are compared to the average response from both reference side patches. All traces reflect percent-change in coherence with respect to the tagging baseline interval, and black traces reflect differences of the corresponding pair. Shaded regions around traces in **B** are 95% bootstrapped CIs (cousineau-corrected), and horizontal bars in **B** as well as stars in **A** reflect significance as per cluster permutation test.

This tagging-baselining procedure enabled the comparison of tags other than just diametrically opposite, frequency-counterbalanced pairs alone. Firstly, this alternative comparison method reproduced our original analysis, showing a significant modulation of the outer cued tag as compared to the outer tag on the reference side (significant cluster from 0.4s-1.2s after cue onset, p = 0.0002), and then later a significant modulation of the middle tag on the cued side compared to its reference (0.57s-0.87s after cue onset, p = 0.011). Additionally, it also showed a numerical (but not significant at the cluster level) attentional boost at fixation immediately after cue onset, which returned back to baseline once attention was allocated to the cued outer location.

Using a jacknife approach (see *“General Statistical Analysis”* in Methods), we determined the peak attentional modulation at fixation relative to the reference side to occur at t = 0.275s after cue onset (95% CI: [0.270, 0.287]s). With the same jackknife approach, we determined that the peak attentional modulation at the cued location relative to the reference side occurred at t = 0.569s after cue onset ([0.562, 0.587]s). These values allowed us to produce some rough estimates regarding the physical parameters of the endogenous shifts of attention that took place during the task. We can estimate that the actual shift of attention took ∼0.3s (the difference between the above mentioned peak timings at fixation and the cued location, see *“Latency comparisons”* in Methods). Given a distance of 6.3dva from fixation to the center of the peripheral section that was cued, this suggests an attentional movement speed of ∼21.5 dva per second. The width of the “middle” locations that we tag here forms 22.22% (1.4dva section width / 6.3dva total distance) of the path between start and end-point. A hypothetical, linearly-travelling attentional spotlight would then directly overlap with the middle location on the cued side for ∼70ms (0.3s * 22.22%). This value is inherently based on assumptions (such as a linearly uniform movement of a hypothetical attentional spotlight), and we are thus limited in the strict conclusions we can draw from this particular duration.

Nonetheless, this value (∼70ms) can allow us to, with an in-silico approach, determine whether attention shifted too fast for us to be able to measure it. Although RIFT has been widely used to track spatial shifts of attention, the shifts being studied with it are usually sustained for at least a few hundred milliseconds (Arora et al., 2025; Zhigalov et al., 2019). Our above estimate of the duration of enhancement (if an attentional spotlight did indeed travel across visual space) is much shorter (∼70ms). Thus, we constructed simulated RIFT-EEG data (Figure 4A-B, see *“Simulation Analysis”* section in Methods for details; parameters were chosen to approximate the middle location tagging responses). This was to determine the minimum duration of an attentional effect that we could have measured, as well as establish a lower bound for the speed of a hypothetical moving spotlight. By parametrically varying attentional gain and duration (the two attentional parameters visualized in Figure 4B) with the simulated data, we computed estimates (Figure 4E) of how strong the attentional gain would have to be at various attentional durations for us to have picked it up with our pipeline.

**Figure 4:**
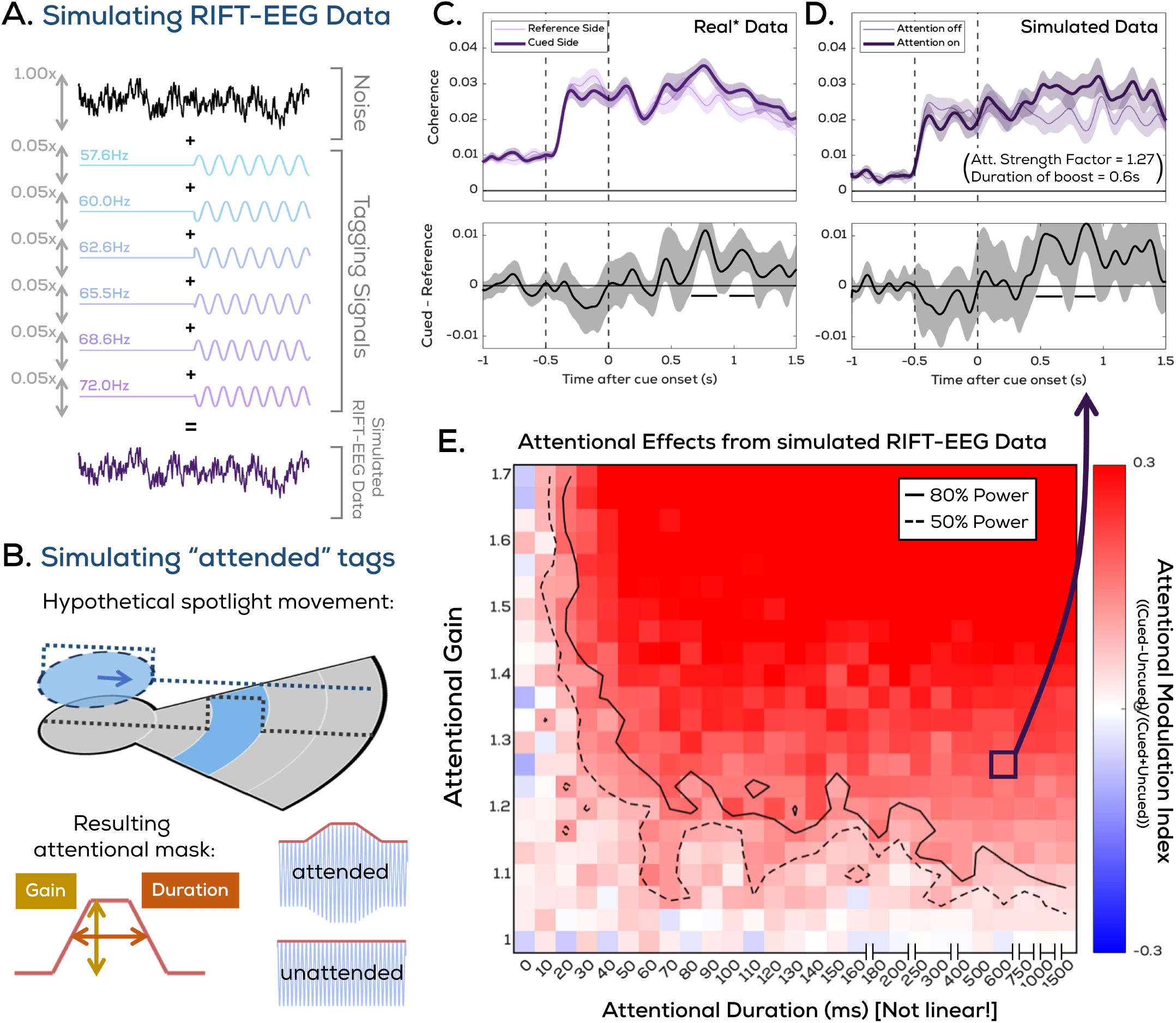
Simulation analysis to assess the limits of RIFT attentional modulation in the current dataset. **A**. We simulated RIFT-EEG data for 100 in-silico participants using pink noise and low-amplitude phase-randomized tagging sinusoids; see *“Simulation Analysis”* in Methods for details. **B**. Then, we simulated “attended” trials using an attentional boost with two variable parameters: gain and duration. We varied these parameters across a range of values (**C-D:** example comparison for a specific parameter set with real attentional modulation observed from collected data), to determine whether RIFT and our analysis pipeline is incapable of measuring short-lived attentional modulations. **E**. For each parameter set, we obtained an average Attentional Modulation Index (AMI) for random samplings of 24 participants, and assessed reliability of the effect by computing what percentage of random samplings showed a significant effect (counter lines in **E**). *See *“Simulation Analysis”* in Methods for details on how the real data traces were prepared for this comparison.

This allowed us to draw a curve showing how long and how strong an attentional modulation would have had to be to achieve a certain level of statistical power (defined as how often our repeated bootstrapping procedures resulted in a significant RIFT modulation). This curve can then be interpreted to reveal whether we would have expected effects of certain durations and gains in our EEG data. If we assume the same level of attentional gain as that which qualitatively matches the real effect (Figure 4C-D) observed at the middle location later in the trial (gain = 1.267), we would be able to observe significant attentional modulations for effects as short as ∼60ms with 80% power. This simulation analysis therefore shows that our experimental approach yielded ample sensitivity to measure an attentional modulation at the intermediate locations, given our data-driven duration estimation of ∼70ms (described above). Thus, either attention does not enhance processing at intermediate locations, or, it does so for less than ∼60ms.

Given that the middle location on the cued side was never task-relevant, it is unclear why attention appears to be allocated to this location after it is allocated to the cued location. Such an invisible probe at intermediate shift locations has not been included in previous studies, so it is also unclear if this is something we would have expected. To confirm whether attention had shifted “normally”, we checked whether two other commonly used markers of spatial shifts of attention showed the pattern that is traditionally expected in our data. These were lateralized differences in alpha power, and small shifts in gaze position (<0.05dva, no saccades were made). Topographies of alpha power following the onset of the cue looked qualitatively different depending on cue direction (Figure 5A), as is known from studies that use these topographical differences to decode attentional locus (Samaha et al., 2016). We explicitly tested for the well-established effect of alpha lateralization (a decrease in alpha power in the contralateral hemisphere following cue onset). Subtracting left cued from right cued trials showed significant differences in alpha power in several channels (Figure 5B), and averaging the left cued minus right cued differences across these significant channels showed a clear effect following cue onset (significant cluster 0.36s-1.5s after cue onset; p < 0.0002). This effect (Figure 5C) was sustained for at least ∼1.5s, which indicated that attention remained lateralized after the onset of RIFT modulations at the middle location.

**Figure 5:**
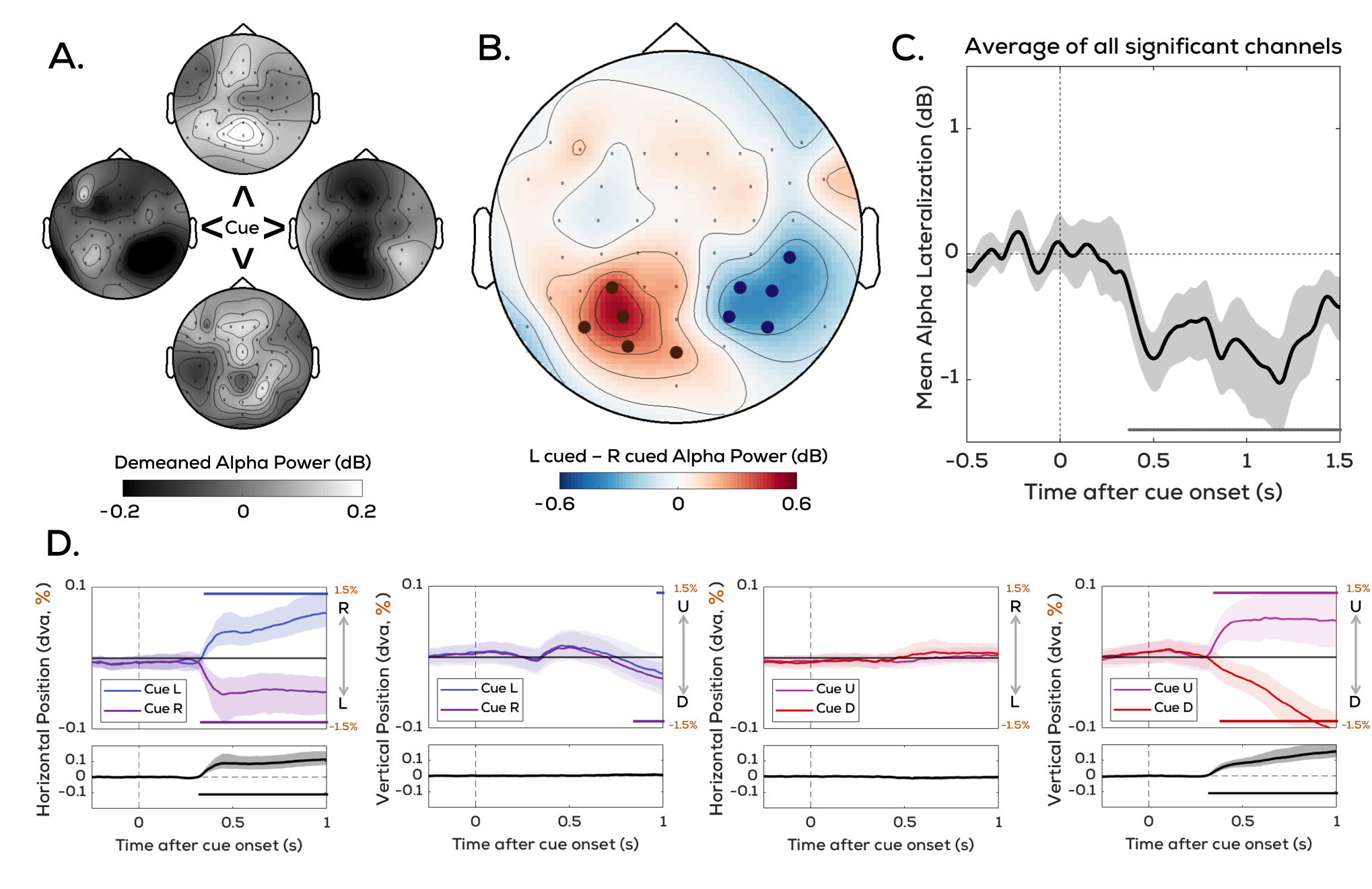
Alpha oscillations and small (<0.05dva) gaze biases confirm covert shifts of attention. **A**. Topographical distributions of alpha oscillations qualitatively differ depending on which direction (right/down/left/up) was cued. **B**. Specifically, alpha lateralization could be tested for between the left and right cued trials, **C**. which over time revealed that alpha power was indeed reduced in the hemisphere contralateral to the cued direction. **D**. The set of four panels represents comparisons in mean gaze position between trials where either left vs. right trials were cued (left two panels) or up vs. down trials were cued (right two panels), based on horizontal gaze changes along the x-dimension or vertical gaze changes along the y-dimension. Marked channels in **B** reflect channels with significant differences. Shaded regions in **C-D** reflect 95% bootstrapped CIs, and horizontal bars reflect significance based on cluster test. In **D**, distances on the y-axis are reflected both in dva and in percentage of the distance from fixation to the center of the cued section of the stimulus.

Similarly, gaze position (Figure 5D) also reflected the cued location, with an amplitude consistent with micro-saccadic gaze activity which is known to track the locus of attentional shifts (Engbert & Kliegl, 2003; Van Ede et al., 2019). Gaze position traces revealed whether the cue pointed up, down, left, or right (up vs. down cued trials, vertical position comparison, significant cluster from 0.33s-1.5s after cue onset, p < 0.0002; left vs. right cued trials, horizontal position comparison, significant cluster from 0.32s-1.5s, p < 0.0002). Thus, standard measures of attentional shifting confirmed that the initial attentional shift had taken place as expected.

## Discussion

We show that during a covert, spatial shift of attention, sensory processing is only facilitated at the target location and not along the trajectory of the shift.

Our results suggest that sensory processing does not “travel” from the start to endpoint of a covert attentional shift. They instead show that sensory processing “teleports”, without any enhancement at intermediate locations. From a functional point of view, this makes sense; it is not necessary that there will be behaviourally relevant visual information between two locations that attention shifts across. This finding does deviate from previous work, as attentional facilitation of locations between an attentional shift has been previously found. However, these cases differed from our design in one of two ways: firstly, a methodological difference in some cases is the use of visible probes to localize attention (Naber et al., 2026; Shulman et al., 1979), in which case the probe itself (or even the awareness of its upcoming appearance) may draw attention and alter how attention is allocated across the visual field. Here, we circumvent this by using RIFT, an invisible tracker of attention, which allows us to assess attentional allocation without any probe-driven effects. Secondly, facilitation at intermediate locations has been found in object-based attention (Jeurissen et al., 2016; Pooresmaeili & Roelfsema, 2014) or curve-tracing designs (Houtkamp et al., 2003; Scholte et al., 2001). In these cases, however, the intermediate locations along the trajectory of attention have some functional relevance to the task at hand. In the case of curve-tracing tasks for example, attention must follow the full path of the curve to determine the corresponding endpoint. In our study, the intermediate location has no connection to the task and comprises no relevant (or even irrelevant) stimuli. This distinction in relevance is most likely what determines whether sensory processing is boosted at intermediate locations during a shift of attention.

With the distinction we establish above in mind, we can also draw a broader conceptual link from our findings on covert attention to the functioning of overt attention. Eye movements primarily involve saccadic motion, where gaze position jumps ballistically from one point to another, with perceptual input being mostly suppressed along its trajectory (Ibbotson & Krekelberg, 2011). However, an alternative to this is smooth pursuit eye movements (Robinson, 1965), where gaze position travels across the visual field to track a moving object. When describing how covert attention affects visual processing at intermediate locations, previous designs with relevant stimuli at these intermediate locations have found an enhanced facilitation there (Houtkamp et al., 2003; Jeurissen et al., 2016; Pooresmaeili & Roelfsema, 2014; Scholte et al., 2001). This can be likened with the case of smooth pursuit eye movements - a gradual, active shift of attention across visual space is warranted in both cases because of the presence of relevant information along the whole shift trajectory. In contrast, our task, where only the endpoint alone is relevant, can be compared to the case of saccadic eye movements, where the endpoint holds the to-be-attended stimuli, and intermediate locations are not worth attending. This account aligns with previous work that has found a reduction in overall sensory sensitivity during covert attentional shifts (Zhang et al., 2026), similar to how our perception is limited during a saccadic eye movement. Our results draw a conceptual parallel between this frequent implementation of covert attention (a simple, voluntary shift of spatial attention between two points) and the most frequent implementation of overt attention.

The RIFT response that we use here stems predominantly from early visual cortex (EVC; Duecker et al., 2025; Ferrante et al., 2023; Minarik et al., 2023). What we measure when using RIFT as a marker of attention is simply whether EVC is more responsive to (imperceptible) stimuli presented at a particular location. However, attention also has well-established effects on more downstream regions of the visual hierarchy (Kastner et al., 1999; Luck et al., 1997). We have also previously shown that attentional signatures may emerge in more downstream parts of the visual hierarchy in the absence of (Arora et al., 2025) or independent of (Arora, Van der Stigchel, et al., 2026) attention-induced RIFT modulations. The fact that we do not observe an effect at intermediate locations here therefore does not discount that representations of intermediate locations during a covert shift may be enhanced elsewhere in the brain. Our findings here simply show that while making a covert shift of attention between two points, we do not boost the processing of stimuli presented at the intermediate location before covert attention arrives at the intended endpoint.

A surprising finding was that the RIFT response from the intermediate locations was enhanced, but only after the covert shift of attention had already “arrived” at the cued location. Firstly, this finding demonstrates that our method does have the sensitivity to measure attentional modulations at these middle locations. Theoretically, though, observing this enhancement after that of the cued location is unexpected, since this middle location was not task-relevant. Had this effect been observed before, or even simultaneously with the effect at the endpoint cued location, we would have interpreted this effect as “travelling” attention. The fact that it occurs afterwards makes it unclear what causes this modulation. The cued location effect also continued to be sustained beyond the onset of the middle location effect, and participants performed the task well (targets were often yet to be presented at the cued location when this intermediate modulation was observed). Thus, this enhancement at the middle location does not reflect the disengagement of attentional resources from the cued location upon task completion. One possibility is that upon shifting attention to the cued location, the “wedge” of the propeller-shaped stimulus is unavoidably registered as an object, due to its salient boundary outlines. This later boost at the intermediate location might then reflect an ongoing spread of attentional facilitation throughout the wedge of the cued location inward from the cued location, as has been proposed and observed for object-based attention tasks (Hollingworth et al., 2012; Jeurissen et al., 2016). Alternatively, it is possible that the variable upcoming target location (targets were presented at a range of angular eccentricities within the cued outer segment) promoted an expansion of the attentional focus, which we measured as increased attentional facilitation at the middle location. Although we cannot conclusively determine what caused this later enhancement here, future work may test these possibilities.

The above results show that several questions about how attention spreads across the visual field still remain open. With our data, we are now able to visualize how attentional modulation evolves over time across the whole visual field (see Supplementary Video 1). By leveraging the temporally resolved, simultaneous, yet invisible nature of RIFT, we visualize attentional shifts here with a level of complexity not previously possible without invasive recordings. This can serve as a starting point for future work into resolving the existing questions surrounding attentional shifts.

Comparing the timings of various attentional modulations across the visual field allowed us to measure the duration of a covert shift of attention. This, in turn, allowed us to estimate the duration with which a hypothetical spotlight of attention would modulate the intermediate location if it were to travel from the start to endpoint in our task. Here, we estimated a duration of ∼70ms to cross a 1.4dva intermediate distance, resulting in an approximate value for the speed of covert attention (∼21.5 dva/s). This is substantially longer than the minimum duration of ∼60ms that we could reliably measure from the RIFT response at the intermediate locations based on our simulation analyses, thus providing further evidence that attention did not travel. In this simulation, we parametrically varied attentional gain and duration. To get a sense of the minimum attentional duration we could have picked up on, we retrieved the value for the other parameter (gain) by taking the strength of the RIFT modulation observed in the real EEG data we collected. For this we use the RIFT modulation at the middle location, since the simulation analysis asks whether the middle location would have shown us an attentional effect. However, the outer location is what was actually cued, and this middle location was never task-relevant, and yielded a slightly weaker attentional modulation than the endpoint. Thus, by taking the attentional modulation at the middle location as a reference for our gain parameter, we may underestimate the actual attentional gain, and thus overestimate slightly the minimum duration of the effect that we could pick up on.

We show here that when participants are cued to a particular direction, even in the absence of any saccadic eye movements, there are small deviations in gaze (<0.05), consistent with established work on microsaccadic eye movements during a shift of attention. A remaining question then is whether tagging modulations truly represent attentional boosts, or simply, tagging flickers becoming closer to the fovea. Our data suggests that the latter is not the case, since we show that attentional boosts at the middle location in the cued direction emerge after attentional boosts at the outer (cued) location. If our tagging modulations were purely driven by gaze position, we would not have expected a delay between these two effects, but rather a simultaneous onset. Furthermore, it has been shown numerous times that attention-driven RIFT modulations and fixational gaze position changes of this small magnitude are not correlated (Arora, Hustá, et al., 2026; Arora, Van der Stigchel, et al., 2026; Arora et al., 2025). Thus, it is highly unlikely that the tagging results we see here are the result of gaze position changes.

## Conclusion

Here, we use a RIFT-EEG task to show that during a covert, spatial shift of attention, sensory processing is not facilitated along the trajectory of the shift. In doing so, we show that preferential processing of stimuli, which is a hallmark signature of attention, is restricted only to the endpoint of the shift, rather than moved across the visual field as per the traditional “spotlight” picture. More broadly, this study forms a first step towards non-invasively tracking attention over the whole visual field to reveal insights in the various domains of visual cognition that are shaped by attention.

## Methods

### Participants

24 healthy participants (age: 22.17 ± 1.99 years old, mean ± std; assigned sex at birth: 17 female, 7 male) with normal or corrected-to-normal vision took part in our study. None reported any history of epilepsy or actively being under medication for psychiatric diagnoses. Participants were compensated either with €20 or an equivalent amount of participation credits as per Utrecht University’s internal participation framework. The study was carried out in accordance with the protocol approved by the Faculty of Social and behavioral Sciences Ethics Committee of Utrecht University.

### Protocol

Participants joined for a 2 hour experimental session, at the start of which they received procedural information and provided informed consent, date of birth, assigned sex at birth, and dominant hand information at the beginning of the session. Once EEG equipment was set up, participants were seated 72cm from the screen with a chinrest. After eye-tracker calibration, the experiment was explained using a visual guide and verbal script. Following these instructions, practice trials were conducted until participants reached approximately 70% accuracy. Once the desired accuracy was achieved, participants began the experiment, which lasted approximately 1 hour (15 blocks of 64 trials each). Eye-tracker calibrations were performed again after every three blocks. Compensation was awarded when applicable, and the session was ended. Data collection was conducted between October 2025 and January 2026.

### Display Apparatus

The experiment was displayed using a ProPixx projector (VPixx Technologies Inc.; resolution = 960 × 540 px, refresh rate = 1440 Hz), a device specialized for the high-frequency display that RIFT necessitates. Rear-projection was used (projected screen size = 49 × 27.6 cm) was used for the experimental display. The task was coded in MATLAB and PsychToolBox (Kleiner et al., 2007).

### Experimental Design and Procedure

Participants performed an orientation discrimination task following an endogenous spatial cue (Figure 1A). The background shape consisted of four angular sectors (combined with a central circular area), hereafter referred to as the “propeller” stimulus. This specific stimulus design was selected to maximize area in the periphery, since higher area provides a higher RIFT tagging response.

Each trial began with a true baseline period of variable duration (0.75-0.80s), during which only the propeller stimulus and a fixation cross were presented. Following this, a tagging baseline period began, in which nothing compared to the true baseline period differed perceptually for the participant, but stimuli began to flicker (see *“Tagging Manipulation”*). Tagging continued until response. After the tagging baseline period (0.5s), an arrow cue (0.1s) was shown that pointed either right, down, left, or up. This cue indicated to participants with 100% validity where the target that required a response from them would be presented. A variable cue-target delay period (0.4-1.3s with respect to cue onset; not uniform, see below) then followed. Finally, a pair of targets was simultaneously presented; one at the cued location, and one at the location diametrically opposite to this. Targets (Landolt-C stimuli; 0.1s) were always presented in the most eccentric section of the propeller stimulus, never in the intermediate sections. The angular eccentricity of targets could vary to place them anywhere within this most eccentric section, but the cued and uncued targets were always diametrically opposite. Participants were instructed to report the orientation of the cued Landolt-C (i.e., whether its opening faced right, down, left, or up) with arrow keys. Feedback was presented in the form of a green check mark or red cross for correct or incorrect trials respectively. If no response was received within 1.3s, or a different button was pressed, the trial ended (mean 2.05% of trials, median 1.98% of trials) and feedback was given as per incorrect trials.

Cue-target delays (SOA; stimulus-onset asynchrony) were not uniformly random throughout this interval, instead, shorter intervals were less likely than longer intervals. This was to ensure that we could still obtain immediate shifts of attention in anticipation of sudden target onsets, without frequent short SOAs preventing us from observing attentional allocation before the target onset. Within the range of target onset intervals (0.4-1.3s after cue onset; continuous, not discrete), the likelihood of a given SOA x followed a probability distribution P(x) = x (as opposed to a uniformly random distribution, P(x) = constant). Essentially, this meant that SOAs around 0.4s almost never occurred, and SOAs around 1.3s had the highest probability, with probabilities gradually increasing from 0 to maximum for intermediate SOAs.

Participants completed 960 trials each (15 blocks x 64 trials per block). The direction in which the cue pointed (right, down, left, or up) and the position of the cued and reference side tags (two configurations) were counterbalanced. The orientation of both landolt-Cs was selected randomly on each trial, as was their angular eccentricity within the area in which they were displayed.

### Tagging Manipulation

Here, we quantified the allocation of attention to various locations on the screen using Rapid Invisible Frequency Tagging (RIFT). This technique involves flickering the luminance of specific patches or stimuli on screen faster than can be perceived, and recording the oscillatory response to these luminance changes as a measure of visual processing (Figure 1B).

In this task, 9 patches across the propeller stimulus with four wedges of segmented arms were simultaneously tagged. Throughout this study, these tags have been labelled according to (i) their relationship with the direction in which the cue points (see Figure 1C), and (ii) their location. With respect to the cue, tags are labelled either as “cued” if they are on the cued side, or “reference” if they are diametrically opposite to the cued side. With respect to position, tags are defined as “outer” when referring to the most peripheral segments of the wedges, resulting in “outer cued side” and “outer reference side” tags (tagged using 57.6Hz and 60.0Hz counterbalanced). Similarly, tags are defined as “middle” when referring to the middle segment of the wedges, resulting in “middle cued side” and “middle reference side” tags (tagged using 62.6Hz and 65.5Hz counterbalanced). The “other” tag covers both remaining wedges that are 90 degrees away from the cued direction - all four of these patches flicker at the same frequency (72.0Hz). Lastly, the central area around the fixation cross is tagged, resulting in a “fixation” tag (68.6Hz). This precise set of six unique tagging frequencies (57.6Hz, 60.0Hz, 62.6Hz, 65.5Hz, 68.6Hz, 72.0Hz) was used because they amount to integer multiples of the 1440Hz refresh rate used here. Temporal precision of the displayed stimuli was ensured using PsychToolBox’s Screen(‘Flip’) command.

### Stimuli

The screen background was maintained at a dark grey (RGB: [100, 100, 100]) throughout the experiment. The main propeller stimulus consisted of four angular sectors combined with a central circular area. Each “wedge” component subtended an angle of 60° at the center, meaning that the angular spacing between two adjacent wedges was 30°. Each wedge section had a radius of 7 dva, which was split up into 5 equal parts to make 4 annular sections each 1.4 dva wide. The innermost part formed a circle (r=1.4 dva) around fixation. The fixation cross (0.2 dva) was presented in black, and the directional arrow cue consisted of two lines (length 1.22 dva) with an angular separation of 53°. Black Landolt-C stimuli had a diameter of 1.07 dva. Target visibility was staircased to maintain an even difficulty throughout the task and across participants. Accuracy was maintained around 70% using a staircase procedure (PsychtoolBox QUEST algorithm (Farell & Pelli, 1999)) with parameter settings *β* = 3.5, Δ = 0.01, *γ* = 0.5.

Tagging involved changing the luminance of the pixels within a particular tagged section from fully black (RGB: [0, 0, 0]) to fully white (RGB: [255, 255, 255]) sinusoidally at the corresponding frequency. Rather than achieving this by drawing annular sectors at specific luminances on each frame (which would increase computation time), we achieved the tagged display by simply tagging rectangular regions and masking these with the propeller stimulus which had transparent regions through which the tagging could be displayed (Figure 6).

**Figure 6:**
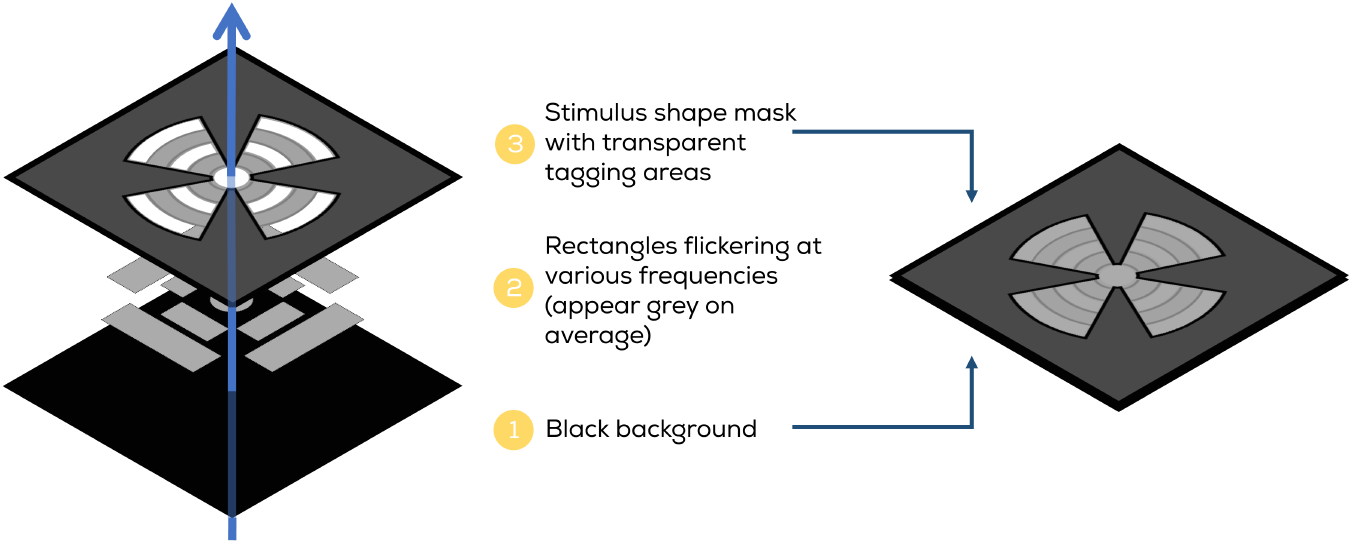
Construction of tagging stimulus. To tag specific patches of the propeller stimulus while still using simple computationally inexpensive drawing functions, we tagged rectangular patches and then masked these with an image corresponding to the stimulus shape.

### EEG recording and pre-processing

EEG data were recorded using a 64-channel ActiveTwo BioSemi system (BioSemi B.V.) at 2048 Hz. Four additional electrodes were placed: two positioned behind the ears on the mastoids, and one each above and on the outer canthus of the left eye. Adequate signal quality from all channels was ensured using BioSemi ActiView software prior to recording. All data analysis was conducted in MATLAB using the Fieldtrip toolbox (Oostenveld et al., 2011). First, EEG data were re-referenced to the average of all channels (excluding poor channels determined by visual inspection, median = 13 [mainly frontal] channels, mean = 14.3 channels). A high-pass filter was applied (0.01 Hz), and then line noise and its harmonics were removed using a DFT filter (50, 100, 150 Hz). Data were then segmented into trials ranging from 1 sec before to 1.75 sec after cue onset. An ICA was performed to remove oculomotor (blink) artifacts, and trials with other artifacts or noise were removed from further analysis as per visual inspection (median = 10.5%, mean = 12.1%). Baseline correction was performed by averaging (and then subtracting from the signal) a window 1-0.5 sec before the cue.

### RIFT Response: Coherence

To get time-varying estimates of the RIFT response strength at the various tagged frequencies, we used a filter-hilbert transform (filter-HT) approach paired with magnitude-squared coherence. Coherence evaluates frequency responses by assessing both the frequency amplitude of the input signal, as well as the degree to which this frequency content is phase-locked over trials. This also means that a set of trials on which coherence is computed gets reduced to one coherence trace (a dimensionless time-varying quantity ranging from 0 to 1); i.e., it is an across-trials measure.

Before computing coherence, the filter-HT approach used here involved first bandpass filtering the EEG data (two-pass Butterworth filter, 4th order, hamming taper), then running a hilbert transform on the filtered data to produce a time-varying magnitude (*M* (*t*)) and phase (*ϕ*(*t*)) estimate per trial. Magnitude-squared coherence is computed between the signal of interest (filter-HT output of the EEG response) and a reference signal; for the latter we use pure sinusoids of the frequency being tested. The magnitude and phase estimates from the filter-HT output and the magnitude and phase of a pure sinusoid at the frequency being tested, were used as per Equation 1 to compute a time-varying coherence measure for a given set of trials, where *x* and *y* subscripts denote EEG data and reference signal features respectively.

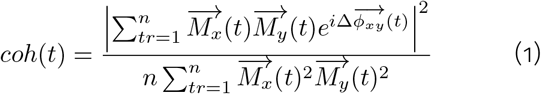

To visualize the tagging-induced spectral peaks with spectrograms, we computed coherence at frequencies from 53.6Hz to 75.2Hz in increments of 0.8Hz using a bandpass filter width of 2Hz. For all further analysis, we computed coherence only at the tagging frequencies, and used a bandpass filter width of 5Hz for increased temporal resolution. See *“Phase-skewing Approach”* for more on how this was done for each frequency.

Coherence is computed for each channel separately. In order to produce spectrograms or traces, a channel-averaging procedure is then conducted. In line with previous studies (Arora, Van der Stigchel, et al., 2026; Arora et al., 2025; Hustá et al., 2025), we selected the top 6 channels with the strongest RIFT response per participant (to allow for small variations in topography across participants) and averaged these to produce a channel-averaged coherence measure. Note that it has been shown that modulations of coherence are independent of the exact number of channels (here six) that are used in this manner (Arora, Hustá, et al., 2026; Arora et al., 2025).

### Phase-skewing Approach

To utilize a high number of unique frequency tags without the limitation of spectral overlap, we implemented the phase-skewing approach described in (Arora, Hustá, et al., 2026). In short, on each trial, we assigned each of the six frequency tags used here a random initial phase. During analysis, we could then take the same set of trials, and skew the EEG responses of each trial by the initial phases of any one frequency at a time, such that only the frequency response of that particular frequency is aligned across trials. Since coherence, the measure used here to assess the RIFT response, measures phase-alignment across trials, this ensured that we could separately measure each of our six tags. Thus, we always ran coherence analyses on the same data six times; once each with the trials phase-aligned to each of the six frequencies used here. This procedure allowed for much improved separability between the tags without sacrificing temporal resolution.

### Simulation Analysis

We used an in-silico approach to obtain additional evidence of whether our frequency analysis methods would have been able to detect short-lived attentional facilitation at the middle tags if there was any. This approach consisted of three overarching steps: 1) modelling RIFT-EEG trial data, 2) modelling the effect of attention onto this data, and 3) mimicking experimental conditions.

#### Generating the Data

We first used pink noise (1/f power spectra) to produce a basic time series signal at 2048Hz reflecting EEG responses. Then, to reflect the RIFT response, we added weighted pure sine waves at each of the six tagging frequencies that were used in this task to the pink noise signal. Naturally, the strength of the RIFT response in this simulated dataset would depend on the first variable parameter: the weight used for the sinusoids as compared to the noise trace. We selected this parameter so as to match the level of coherence resulting from the simulated data as best as possible to the coherence traces obtained from real data. The amplitude of the sinusoidal traces in this case was only ∼1.5% the amplitude of the pink noise traces (Figure 4A). Note that these simulated tags were also randomized in phase; they underwent the same steps described in the “Phase-skewing approach” section to maintain the same ability to separate distinct tags as that with the real data. This weighted summation of pink noise and tagging sinusoids formed a basic EEG-RIFT response used for the simulation analysis.

#### Replicating Attention

Next, we implemented “attention”, the effect of which we later parametrically varied in Figure 4E. Attention was implemented as amplitude gain of the “attended” tag (i.e., scalar multiplication of the corresponding sinusoid by the gain factor – a gain of 1 implies no attentional boost). This produced two parameters; firstly, amplitude gain, the strength of the attentional boost. Secondly, the duration for which this amplitude gain persisted. This amplitude gain was not modelled with an immediate onset and offset in time. Rather, the gain profile over time was obtained by mimicking the effect of a travelling attentional spotlight across the middle tagged area. Specifically, we assumed a uniform attentional spotlight with the same size as the fixation area (see Figure 4B for a visual of the resulting attentional gain mask).

Lastly, to make qualitative comparisons between the cueing effects on real RIFT responses and simulated data, we introduced experimental conditions surrounding trials and participants. We simulated 100 “participants”, each of which had an additional noise multiplier selected uniformly randomly in the range 1 to 1.3. This qualitatively mimicked across-participant variation in the strength of the RIFT response and EEG noise. For each of these participants, we ran 500 “trials”, of which half did not have any attentional boosts (reference RIFT response) and half did (cued RIFT response). We then ran coherence analyses on this data with the exact same parameters as those used for the real data. The only factors that we parametrically varied were those specific to the model’s implementation of attention, specifically, the attentional gain (varied from 1.0 to 1.7 in steps of 0.33), and attentional duration (varied from 0ms to 160ms in steps of 10ms, then at various specific durations until 1500ms).

#### Quantifying outcomes

To concisely present the outcomes of how the strength of attentional effects in the model was impacted by varying attentional gain and duration, we reduced each pair of parameter values to one distribution. For each pair of parameter values, we prepared 1000 subsamplings of 24 participants each (to reflect how many participants we actually recorded from in our EEG experiment). For each subsampling, we averaged the coherence traces corresponding to attended (cued) and non-attended (reference) conditions, similar to how we treated the real data in Figure 2. Then, we computed an Attentional Modulation Index (AMI; computed as (attended - unattended)/(attended + unattended)), and tested whether AMIs were significantly above zero for each subsampling. We recorded the proportion of subsamplings for which AMIs were significantly above zero, and used the contour() function in MATLAB to draw contour lines reflecting 50% and 80% power (% of subsamplings that showed a significant response)

Lastly, when drawing a comparison between the coherence traces of real data and simulated data, we also wanted to qualitatively try to match variance across the two. However, the real traces presented in Figure 2 are averaged over 4 different location conditions, which results in a less variable set of coherence traces across participants. To account for the fact that the model does not observe different location conditions and perform such an averaging procedure, when preparing the coherence traces of real data to compare to the simulated data in Figure 4C, we selected one random location trace per participant. By selecting one random location per participant, we reflect the variance that can be more appropriately matched with the simulated data.

### Alpha-Band Acitivity

Topographical differences in alpha-band (∼10Hz) activity are a common hallmark of spatial changes in attention. We computed time-frequency representations in the 8-13.5Hz range using the `ft_freqanalysis` function in the Field-trip toolbox (Oostenveld et al., 2011). This was done in increments of 0.2Hz for every channel using 3-cycle Morlet wavelets and baselined (dB; 10*log10(signal/baseline)) with respect to a window 0.7-0.2 seconds before cue onset. This range (8-13.5Hz) was then averaged, providing a measure of alpha-band activity relative to the baseline period. To compute alpha lateralization, the alpha-band activity of right-cued trials was subtracted from left-cued trials per participant. To visualize the topographical spread of alpha-band activity across all four cue directions, we demeaned these topographies by subtracting the overall mean from all four cue direction conditions.

### Eye-tracking recording and analysis

An eye-tracker (Eyelink SR; SR Research) was used to record gaze position throughout the experiment at 500Hz. Following an initial 9-point calibration at the start of the experiment, calibration was conducted again after every third experimental block. Both eyes were tracked and later averaged together.

Gaze position data was segmented into trials from 1s before to 1.5s after cue onset. Blink correction was carried out using custom code based on existing work (Hershman et al., 2018). All steps below were carried out separately for x and y position, and separately for both eyes. First, we baseline-corrected position data with an interval 900ms to 200ms before cue onset. We then determined which trials to exclude based both on inadequate fixation, (which was determined by computing whether gaze was further than 1.5 dva away from center for atleast 50ms), and the baseline gaze position being >2 standard deviations away from the mean (% trials removed: mean = 11.72%, median = 10.83%). Lastly, we applied a 10ms uniform smoothing filter to the baselined position data, and averaged both eyes together.

### General Statistical Analysis

Several different analyses produced one difference trace per participant. This includes the coherence results, both absolute and percent change (one coherence trace per participant for each of the two conditions being compared to each other, which were subtracted to produce one difference trace per participant), the gaze data (either a left vs. right cued or up vs. down cued difference trace per participant), and the alpha lateralization data (one alpha lateralization trace per participant being compared to zero). In all these cases, difference traces were tested for significance using a cluster permutation test (Maris & Oostenveld, 2007). With a given set of difference traces, the steps of this permutation test were as follows: firstly, a one-sample t-test was run for each individual timepoint to detect points on the trace significantly different from 0 (p < 0.1). From these values, any clusters of consecutively significant timepoints (minimum 10ms long) were selected, and a cluster t-mass was obtained for each of these by summing the t-values of each point in the cluster. This cluster t-mass was tested for significance by comparing it to a bootstrapped distribution of cluster t-masses. To produce this bootstrapped distribution, we ran 5000 iterations where we randomly flipped the sign of each individual difference trace (so as to not impact the temporal correlation of the traces by shuffling timepoints), and added the highest cluster t-mass to the distribution. Lastly, a cluster from the original difference traces was labelled as significant when its t-mass was in the bottom or top 2.5% of cluster t-masses within the bootstrapped distribution.

### Latency Comparisons

When computing the main cueing RIFT effects at the outer and middle locations, we also tested for differences between their onset latencies. Since the clusters produced by cluster permutation tests are not themselves a direct measure of onset latency (Sassenhagen & Draschkow, 2019), we could not directly compare the onsets of the significant clusters for this. Instead, we built a bootstrapped distribution of latency by running 100 iterations of the outer cued - outer reference and middle cued - middle reference differences, each time sampling 24 participants with replacement from the total set of 24. For each of these samples, we stored the onset of the effect in one of two ways. Either we simply stored the first significant timepoint as per the cluster test (in Figure 2C; this is less susceptible to noise, but the cluster test is not a perfect measure of onset latency), or, simply the onset of the first 2, 5, or 10 consecutive timepoints (in Supplementary Figure 1; this is noisier, but not dependent on timepoints later in the trace as the cluster test is). By showing that both these alternatives converge to the same outcome, we can reliably trust the order of effects suggested by this procedure. This produced a distribution of effect latencies for both the outer and middle location cueing effects, which we compared statistically using a bootstrapped distribution of mean differences.

For determining the peak latency of the attentional modulation at fixation and the cued location as compared to the reference side (Figure 3), we use a jackknife-based approach (Miller et al., 2009). Participant-wise traces can be noisy, and thus selecting the peak timepoint per participant could produce unreliable estimates. On the other hand, simply selecting the peak value from the participant-averaged trace only produces one singular estimate. To balance across these difficulties, we computed 24 participant-wise averages of 23 participants each, by leaving each of the 24 participants once in one of these averages. We selected the timepoint of the peak difference modulation for each of these averages. This set of 24 peak latency estimates could be converted into a mean and a bootstrapped 95% confidence interval that we report as the timepoint at which the effect strength peaked.

## Acknowledgements

The authors would like to thank Joel Mekking, Ruben Hermsen, and Dirk van Tilburg for their assistance with data collection.

## Author Contributions

Conceptualization, K.A., S.C., S.G., S.V.d.S., L.K.; Data Curation, K.A.; Formal Analysis, K.A., S.C.; Funding Acquisition, S.V.d.S.; Investigation, K.A.; Methodology, K.A., S.C., S.G.; Project Administration, K.A., S.C.; Resources, S.V.d.S.; Software, K.A., S.C.; Supervision, S.C., S.G., S.V.d.S., L.K.; Visualization, K.A.; Writing – Original Draft Preparation, K.A., S.C., S.G.; Writing – Review & Editing, K.A., S.C., S.G., S.V.d.S, L.K., M.N.

## Data and Code Availability

Raw and processed EEG data are publicly available at https://osf.io/npwjk/overview.

## Declaration of Interests

The authors declare no competing interests.

## Supplementary Material

**Supplementary Figure 1:**
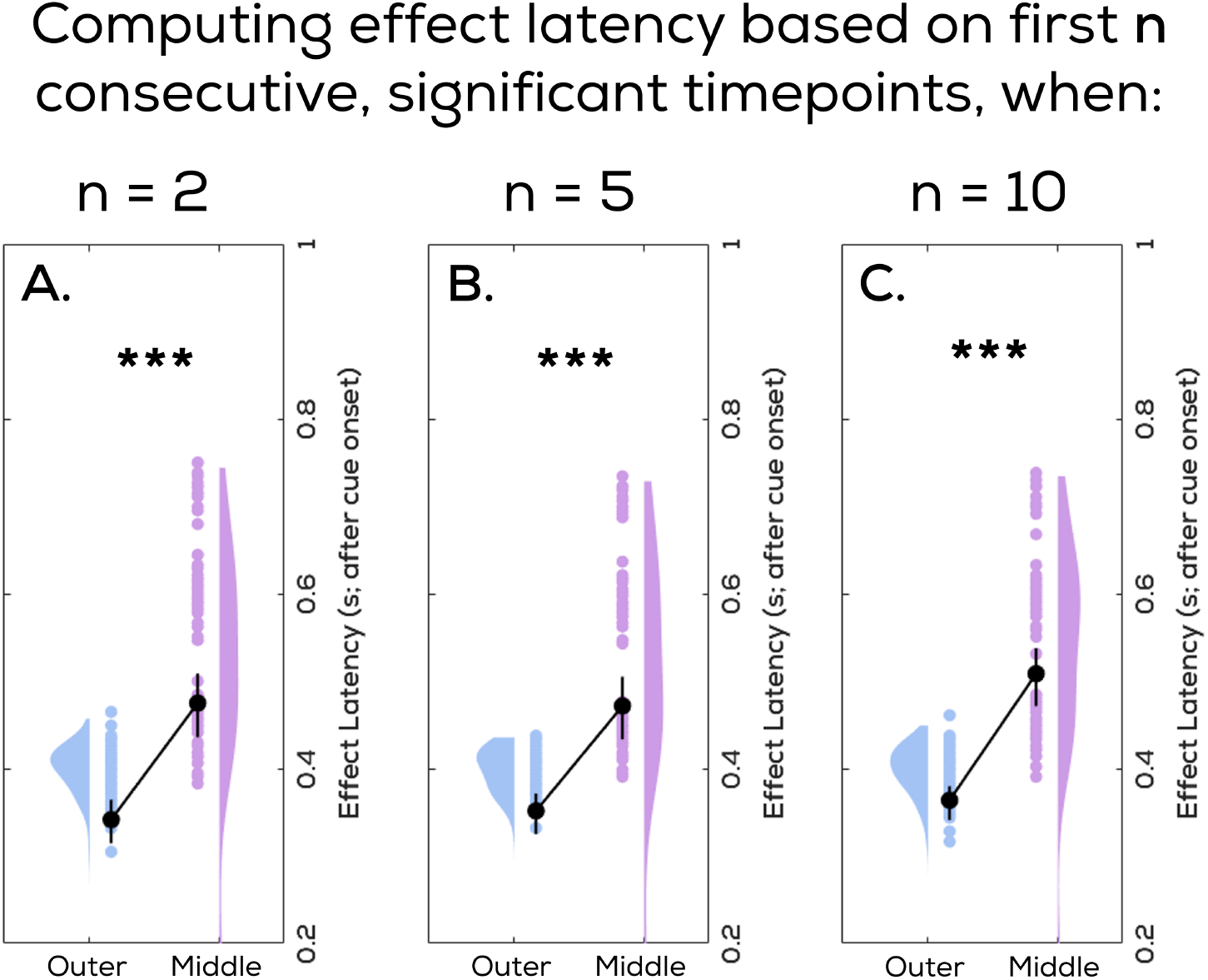
Outer tag modulation precedes middle tag modulation even with alternative latency measure. In the manuscript, we compare the latencies of the attentional modulations to the RIFT responses from the outer and middle locations using bootstrapped analysis based on the first timepoints of significant clusters. Here, we show that the results from that version also hold when, rather than using the first timepoints of significant clusters, we instead use the first few consecutive significant timepoints, regardless of the outcome of the permutation cluster test. Panels **A, B**, and **C** show comparisons where onset latency is defined as the first moment where 2, 5, or 10 consecutive timepoints respectively are significant on a paired-samples t-test (alpha = 0.05). All comparisons show significant results (p < 0.01 in all cases).

**Supplementary Video 1:** Using RIFT, we are able to visualize the distribution of attention over time as it progresses through a shift of attention from the center to the periphery of the display stimulus. We produce these results as a video over time. See https://drive.google.com/file/d/1a2QWIi4SCXinvpndJe5zpncvjSgbCMUY/view

## Notes

https://osf.io/npwjk/overview

### Competing Interest Statement

The authors have declared no competing interest.

